# Dynamic metabolic and molecular changes during seasonal shrinking in *Sorex araneus*

**DOI:** 10.1101/2023.10.02.560485

**Authors:** William R. Thomas, Cecilia Baldoni, Yuanyuan Zeng, David Carlson, Julie Holm-Jacobsen, Marion Muturi, Dominik von Elverfeldt, John Nieland, Dina K. N. Dechmann, Angelique Corthals, Liliana M. Dávalos

## Abstract

To meet the challenge of wintering in place many high-latitude small mammals reduce energy demands through hibernation. In contrast, short-lived Eurasian common shrews, *Sorex araneus*, remain active and shrink, including energy-intensive organs in winter, regrowing in spring in an evolved strategy called Dehnel’s phenomenon. How this size change is linked to metabolic and regulatory changes to sustain their high metabolism is unknown. We analyzed metabolic, proteomic, and gene expression profiles spanning the entirety of Dehnel’s seasonal cycle in wild shrews. We show regulatory changes to oxidative phosphorylation and increased fatty acid metabolism during autumn-to-winter shrinkage, as previously found in hibernating species. But in shrews we also found upregulated winter expression of genes involved in gluconeogenesis: the biosynthesis of glucose from non-carbohydrate substrates. Co-expression models revealed changes in size and metabolic gene expression interconnect via FOXO signaling, whose overexpression reduces size and extends lifespan in many model organisms. We propose that while shifts in gluconeogenesis meet the challenge posed by high metabolic rate and active winter lifestyle, FOXO signaling is central to Dehnel’s phenomenon, with spring downregulation limiting lifespan in these shrews.

## Introduction

Wintering adaptations vary widely across animals and the most common strategies are conserving energy by lowering metabolic rates and activity (e.g. hibernation or torpor) or migration. By contrast, a handful of species that neither migrate nor slow their metabolism have evolved a different wintering size plasticity known as Dehnel’s phenomenon (1–4). This pattern of growth, best studied in the Eurasian common shrew, *Sorex araneus*, is unlike that of almost every other mammal. Instead of growing continuously to adulthood, juvenile *S. araneus* grow to an initial maximum size in the first summer of their disproportionately brief life (∼1 year). In anticipation of harsh winter conditions, shrews then shrink, reaching a nadir in winter, followed by rapid regrowth to their adult, breeding size in spring. This plasticity is not just a retooling of overall body size, but also occurs in many vital organs, including the liver, brain, and spleen (2,5). Although Dehnel’s phenomenon has garnered much interest because of its potential for regenerating organs (6), metabolic and regulatory changes during this wintering adaptation remain largely unknown.

Dehnel’s phenomenon corresponds to the unique physiological constraints shrews face. *S. araneus* have one of the highest mammalian metabolic rates measured to date (7), requiring high and constant food intake throughout the year (8). But supplying energy to maintain this high metabolic rate is especially difficult in autumn and winter, when temperatures decrease and resources become scarce (9,10). Therefore, Dehnel’s phenomenon is construed as an evolutionary adaptation that compensates for continuously high energetic demands by reducing body and organ size, particularly of energy-expensive tissue (11). In support of this hypothesis, absolute resting metabolic rate decreases with shrew size in winter, despite harsh extrinsic conditions (10,12,13)

Wintering strategies encompass a spectrum of related traits and physiological processes (14). Changes in other wintering strategies, such as metabolic shifts observed in hibernation (15,16), may thus share common mechanisms with those involved in Dehnel’s phenomenon. For example, mechanistic analyses of mammalian hibernation have identified a toolkit of genes that regulates vast metabolic changes (15–17), including those associated with increased fatty acid oxidation simultaneous (18), as hibernators rely on gradual fat utilization over winter. Research on shrew physiology indicates they also rely extensively on lipid metabolism in winter (8) and may share at least some molecular mechanisms with hibernation.

But *S. araneus* remain active throughout winter, undergoing drastic shrinkage while navigating a metabolically demanding environment. This active wintering strategy contrasts with hibernation, suggesting key differences in metabolic and regulatory processes. For example, wintering shrews show rapid fat turnover, with 50% fat turnover rates decreasing from 4.5 hours in summer juveniles to 2.5 hours in winter (8). Consequently, metabolic profiles may differ significantly between these strategies. In hibernators there is also a downregulation of genes involved in glucose breakdown (18). In general, when energy stores are depleted, glucose is produced from the liver through gluconeogenesis, which is then released into the bloodstream and metabolized in the mitochondria via oxidative phosphorylation (19). Unlike hibernators that suppress this response, wintering shrews seem to increase this starvation response as environments become harsher (10). Thus, we hypothesized that shrews shrink by upregulating fatty acid metabolism, with potential similarities to hibernators, but unlike them supplement their energy budget with glucose to maintain their active winter lifestyle.

To examine metabolic dynamics and regulatory changes throughout Dehnel’s phenomenon, and its potential parallels to other wintering strategies, we collected phenotypic, metabolomic, transcriptomic, and proteomic data from a shrew population across seasons matching the size plasticity. We focused on blood, which circulates metabolites across the organisms, and the liver, where many functions that regulate metabolism take place, including lipid and sugar processing. Our results reveal seasonal changes in lipid metabolism and gluconeogenesis during shrew size plasticity, which appears to relate to mitochondrial function. Metabolic and size changes are regulated by pathways that include FOXO and insulin signaling, yet incur a cost to lifespan. By exploring these mechanisms, we offer novel insights into the interplay between metabolism, size, and longevity, shedding light on both the wintering strategy of the common shrew and broader mammalian physiology.

## Results

### Dehnel’s phenomenon in German shrew population

Comparisons of body mass throughout the cycle confirmed patterns of size changes (Fig 1A). Analyzing the 24 shrews collected for all stages of Dehnel’s phenomenon (large summer juvenile=5; shrinking autumn juvenile n=4; small winter juvenile n=5; regrowing spring adult n=5; regrown Summer Adult n=5), we found body mass shrank as expected between summer juveniles and winter juveniles (BM_Wij-Suj_=-1.00 g, p_Wij-Suj_<0.028), with regrowth as they matured to spring adults (BM_SpA-Wij_ =3.724 g, p_SpA-Wij_ <0.0001). Shrew liver mass reached a minimum as autumn juveniles (liM_Auj-Suj_ =-904.1 mg, p_Auj-Suj_=0.0949), with the significant portion of regrowth occurring between the winter juvenile and spring adult phases (liM_SpA-Auj_=536.7 mg, p_SpA-Wij_ <0.001). These size changes both validated previous research and were used to guide metabolomic, transcriptomic, and proteomics characterization of Dehnel’s phenomenon.

**Fig 1.**
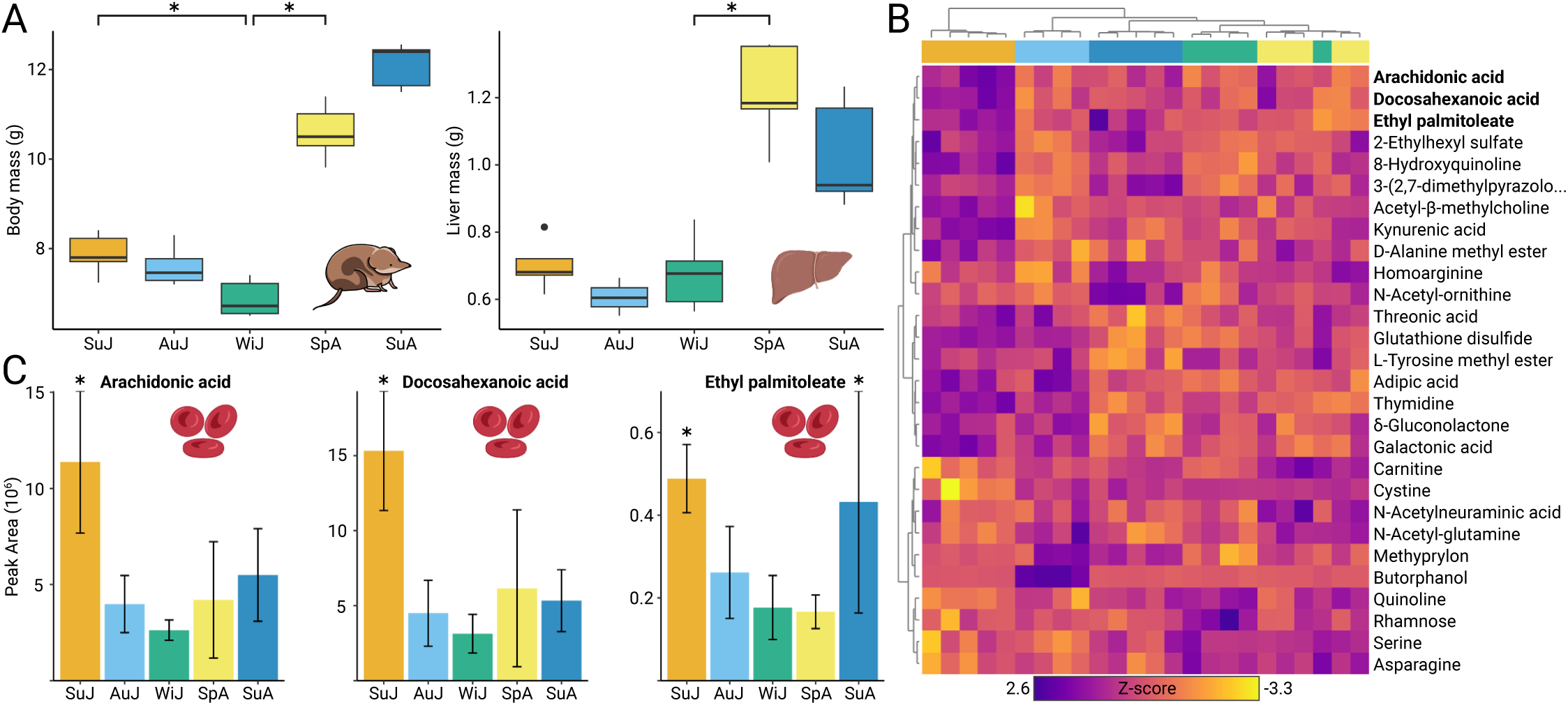
Confirming Dehnel’s phenomenon and metabolic profiling of the blood metabolome. **(A)** Mass change of body and liver through a year of Dehnel’s phenomenon. Both the body and the liver begin to decrease in mass as autumn juveniles, with body mass at a minimum as winter juveniles, proceeded by regrowth as spring adults. Asterisks represent significant size changes (adjusted p<0.05). **(B)** Heatmap of 28 statistically significant differentially concentrated metabolites between stages of Dehnel’s phenomenon. Hierarchical clustering using these significant metabolites groups each profile into each season. **(C)** Three of these metabolites were lipid metabolites (ethyl palmitoleate, docosahexanoic acid, arachidonic acid), with decreases in autumn, winter, and spring individuals.

### Metabolomic shifts in lipid metabolite concentrations

We characterized the blood plasma metabolome to quantify metabolites throughout the cycle and evaluated the potential shifts in metabolism during shrinkage and regrowth. We validated 250 metabolites found in all seasons with multiple detection methods. Of these metabolites, we found 28 with significant differences in mean concentration across the stages of Dehnel’s phenomenon (p<0.05, Fig 1B). Among differentially concentrated metabolites, three were lipid metabolites; arachidonic acid (AA, p=0.04, F=5.78), ethyl palmitoleate (EP, p=0.02, F=6.79), and docosahexanoic acid (DHA, p=0.04, F=5.86). These three lipid metabolites showed significant decreases in peak area, a proxy for concentration, between summer juveniles and other stages of size change (autumn juveniles, winter juveniles, spring adults), especially in winter juveniles (AA_Wij-Suj_ =-8.7 x 10^7^ peak area, DHA_Wij-Suj_ =-1.2 x 10^8^ peak area, EP =-3.1 x 10^6^ peak area), for which the concentrations of DHA and AA were at their minimum.

### Regulatory changes associated with metabolic shifts

Results showed many differentially expressed genes in the liver between summer juveniles and winter juveniles (shrinking stage), with an upregulation of genes related to mitochondrial, lipid, and glucose metabolism. Comparisons revealed 964 differentially expressed genes (p_adj_<0.05), with 497 significantly upregulated and 467 significantly downregulated in winter juveniles (Fig 2A). We then used a ranked gene set enrichment test to identify functional effects of RNA regulatory change. This analysis identified 25 pathways enriched with differentially expressed genes (21 upregulated and 4 downregulated) (Fig 2D). Oxidative phosphorylation was the most enriched pathway (p_adj_<0.001, Normalized Enrichment Score=2.34, [NES]) consisting of 46 genes. This gene set included 21 upregulated genes that encode mitochondrial complex I subunit proteins (NDUFs) and 19 upregulated genes encoding ATPases and synthases. Other pathways enriched with upregulated genes included glycolysis and gluconeogenesis (p_adj_<0.001, NES=2.09), which contained the gene encoding for glucose-6 phosphatase catalytic subunit 1, *G6PC1* (p_adj_=0.41, Log Fold Change=0.30, [LFC]), and fatty acid metabolism (p_adj_<0.01, NES=1.81).

**Fig 2.**
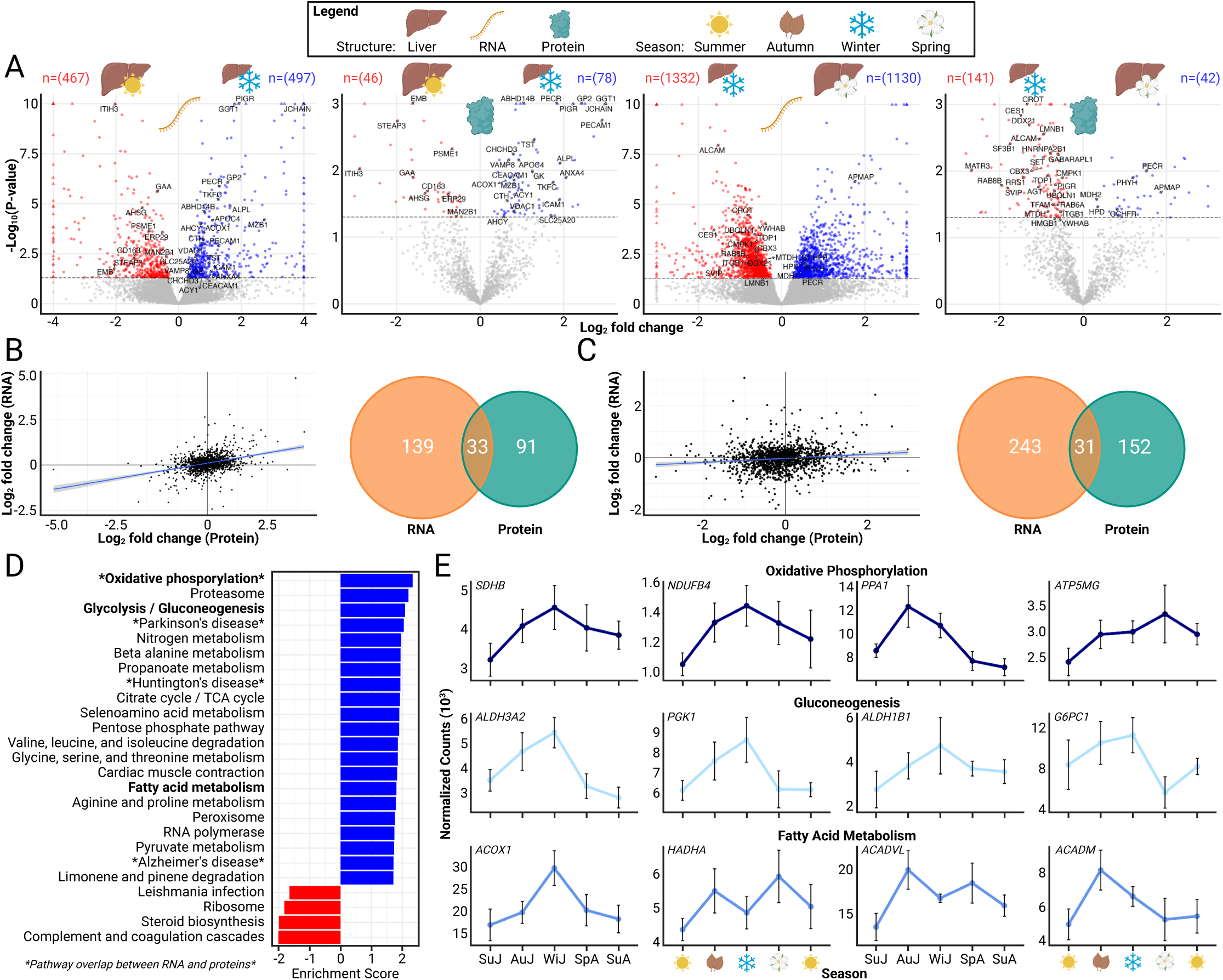
Seasonal regulatory (transcriptomics and proteomics) change in the shrew liver. **(A)** Volcano plots of significant (adjusted p<0.05) differentially expressed genes and proteins (colored) between (from left to right); summer juveniles vs. winter juveniles (shrunk/RNA), summer juveniles vs winter juveniles (shrunk/protein), winter juveniles vs spring adults (regrowth/RNA), and winter juveniles vs spring adults (regrowth/protein) plotted by log fold-change. Triangles represent lower p-values (-log(p)>10) or log fold changes (abs(LFC)>4) beyond the limits of the graph**. (B)** Correlations of log fold change and significant gene overlap between RNA and protein indicate high association between the summer vs. winter juveniles analyses, and **(C)** winter juveniles vs spring adults. **(D)** Pathway enrichment of summer vs winter juveniles between RNA (shown) and protein analyses, both with the largest enrichment of oxidative phosphorylation pathways. **(E)** Normalized gene counts in pathways of interest.

We validated changes in metabolism-related gene expression between summer juveniles and winter juveniles (shrinking stage) with liver proteomics. Despite the liver proteomics (n=1,377 proteins) data set being smaller than the gene expression set (n=24,205 loci), there was substantial overlap in direction, significance, and pathway enrichment of changes between the two. Using a linear model to estimate the correlation between differential change (summer juveniles vs winter juveniles) in gene expression and protein concentration, we found that LFC in gene expression was significantly associated with LFC in protein concentration (F=197.5, df=1373, p<0.001), but the model only explains a small fraction of the variation between them (R^2^=0.13) (Fig 2B). Analyzing differential concentration of proteins, we found 124 proteins significantly changed (p_adj_<0.05) between summer juveniles and winter juveniles (78 increased and 46 decreased in winter) (Fig 2A). Thirty-four of these proteins overlapped with differentially expressed genes, all but one with matching mRNA and protein direction.

Ranked set enrichment for proteomics identified five enriched pathways (p_adj_<0.05), four of which overlapped the RNA pathway enrichment (oxidative phosphorylation, Parkinson’s disease, Alzheimer’s disease, Huntington’s disease). Oxidative phosphorylation was the most enriched pathway in both mRNA and protein expression and included 17 overlapping genes/proteins, 11 of which were found to be significant in at least one of the data sets (*ATP5MG*, *NDUFB4*, *PPA1*, *SDHB*, NDUFV3, NDUFA6, NDUFB8, NDUFS3, ATP5PO, NDUFC2, NDUFS6; all in the same direction). Proteomic analysis did not reveal upregulation in glycolysis/gluconeogenesis (p_adj_=1.00, NES=0.66) pathway, though all but one overlapping gene also increased in concentration in proteomics, including three aldehyde dehydrogenase genes (*ALDH3A2*, p_adj_<0.01, LFC=0.64; *ALDH1B1*, p_adj_<0.01, LFC=0.81; *ALDH9A1*, p_adj_=0.47, LFC=0.35) (Fig 2E). Similarly, proteomic set enrichment analysis did not validate fatty acid metabolism enrichment (p_adj_=0.91, NES=1.00), but there was overlap (11 of 18 genes), including genes that encode proteins that transport (*ACADM*, p_adj_=0.17, LFC=0.40; *ACADVL*, p_adj_=0.26, LFC=0.35) and breakdown of long fatty acid chains (*ACOX1*, p_adj_<0.001, LFC=0.81).

Comparisons between winter juveniles and spring adults (growth) showed extensive change in gene expression coincident with metabolic change, but correlation with proteomics was weak. Of 2,462 significantly differentially expressed genes, 1,130 were upregulated and the rest downregulated in spring adults (Fig 2A). Yet, differential mRNA expression only significantly enriched two pathways, including the downregulation of cell cycle (p_adj_<0.05, NES=-1.77) and proteosome (p_adj_<0.05, NES=-1.91) pathways. Comparing protein concentration between winter juveniles and spring adults (growth), 183 proteins showed significant change (42 increased and 141 decreased) (Fig 2A). While 41 of these overlapped with significant differentially expressed genes, nearly a quarter of them (10) were in the opposite direction. This is unsurprising, as the model between protein LFC and mRNA LFC does not explain a large portion of the variation (*R*^2^=0.02, F=22.73, df=1374, p<0.001) (Fig 2C). Ranked protein enrichment corroborated minimal correlation, as none of the 6 significant pathways: starch and sucrose metabolism (p_adj_<0.01, NES=2.33), TCA cycle (p_adj_<0.01, NES=2.28), cysteine and methionine metabolism (p_adj_<0.05, NES=2.13), valine, leucine, and isoleucine degradation (p_adj_<0.05, NES=1.92), spliceosome (p_adj_<0.05, NES=-1.71), and cell adhesion molecules (p_adj_<0.05, NES=-1.75), overlapped with mRNA gene set analyses. There were, however, individual genes validated in change of direction with proteomic data found in upregulated processes in winter juveniles that decreased in spring adults (LFC<0): oxidative phosphorylation (*NDUFB4*, *PPA1*), gluconeogenesis (*ALDH3A2*, *PGK1*, *ALDH1B1*), and fatty acid metabolism (*ACOX1*).

### Weighted Gene Correlation Network

Differential expression analyses focused on genes of high effect between two size extremes, but large modules comprising genes of *small* effect could also play pivotal roles in regulating metabolic changes across all seasons. We characterized interactions in expression among genes and identified modules of coexpressed liver genes correlated with liver size using weighted gene co-expression network analysis (WGCNA). We identified 37 modules (Fig 3), 10 of which were significantly associated with liver size (p<0.05). Among these, the “grey60” module is a small gene network most positively correlated with larger liver size (n=352 genes, p<0.001, r=0.81, df=21) and the “red” module is a large network positively correlated with small liver size (n=3,268 gene, p=0.001, r=-0.63, df=21). Although no KEGG pathways were significant after multiple test correction when testing for gene set enrichment from the “grey60” module significant genes (n=342 genes, 10 genes were not significantly associated to network), metabolic pathways was highly significant (p<0.01). Thirty-four KEGG pathways from the “red” module significant genes (n=2,706 genes) were significant after multiple test correction (p_adj_<0.05). These pathways included FOXO signaling (p_adj_<0.01), as well as pathways upstream (MAPK signaling, p_adj_<0.05) and downstream (insulin resistance, p_adj_<0.01; longevity regulating pathways, p_adj_<0.05) of this pathway.

**Fig 3.**
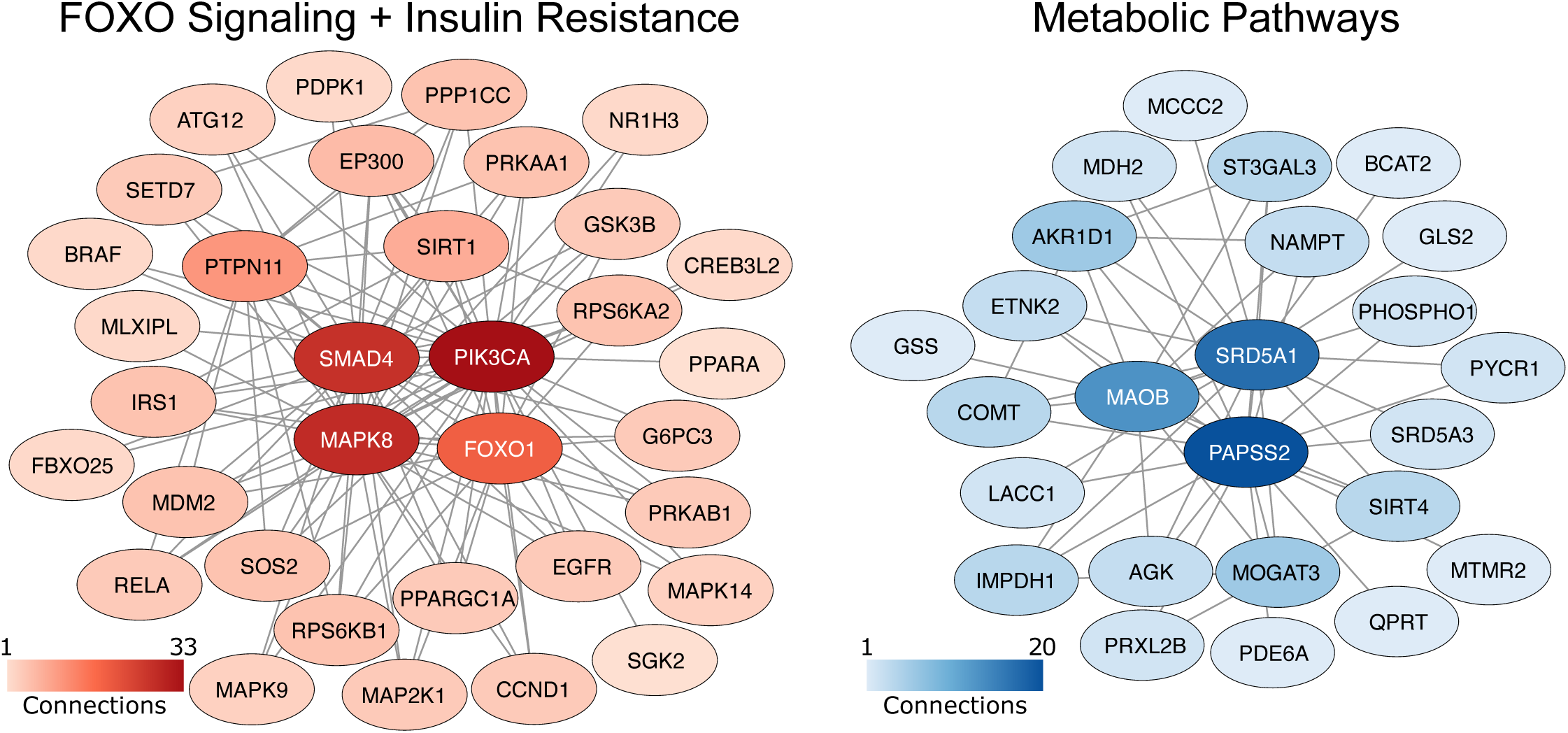
Gene networks highly correlated with seasonal liver size plasticity. Pathway enrichment of the large (n=3268 genes) “red” module highlighted functional associations to FOXO signaling and insulin resistance (plotted genes), while the small (n=342 genes) “grey60” module related to metabolic pathways. Central to FOXO signaling and insulin resistance pathways is the FOXO1 gene (n=19 connections).

## Discussion

Eurasian common shrews, *Sorex araneus*, require constant food intake to sustain their extraordinarily metabolic rates throughout the year (7). Maintaining their energetic budget is particularly difficult in winter, as temperatures drop, as does available food. Within a German shrew population, we found clear signs of a seasonal size plasticity known as Dehnel’s phenomenon (1–3). These shrews radically decrease their size in autumn through winter, with recovery in the ensuing spring. Using this population with confirmed Dehnel’s phenomenon, we could then identify seasonal changes to metabolism and underlying regulatory processes.

We hypothesized that, as in hibernators, wintering relies on fat utilization to maintain metabolic homeostasis in winter, but differs as shrews rapidly turnover fat (8) during shrinkage.

Transcriptomic analyses of winter juvenile shrews revealed upregulation of key lipid metabolism genes, including those involved in fatty acid oxidation: acyl-CoA dehydrogenase medium chain (*ACADM*), acyl-CoA dehydrogenase very long chain (*ACADVL*), and acyl-CoA oxidase 1 (*ACOX1*). *ACOX1*, which encodes the rate-limiting enzyme for oxidizing very long chain fatty acids, is critical for inter-organ metabolic communication, such that liver-specific knockouts of this gene in mice increase adipose tissue browning and circulating levels of polyunsaturated omega-3 fatty acids (20). In shrunken winter juveniles we observe the same correlation, an increase in *ACOX1* expression paired with a decrease in circulating levels of polyunsaturated fatty acids (DHA and AA). This contrasts with hibernators: circulating DHA and AA rise significantly in wintering bears and hibernating arctic ground squirrels (16,21–23). Regulatory similarities and metabolomic differences indicate both wintering strategies rely upon lipid metabolism in the liver. But while hibernators conserve fat reserves over winter, shrews exhibit rapid fat turnover (8), decreasing levels of polyunsaturated fatty acids as they maintain activity in winter.

Regulatory changes related to mitochondrial function further highlight differences in metabolic processes between Dehnel’s phenomenon and hibernation. The strongest signal in both RNA and protein expression was the upregulation of oxidative phosphorylation pathways in winter juveniles, including 21 genes encoding mitochondrial complex I subunit proteins (NDUFs) and the succinate dehydrogenase gene, *SDHB*. Hibernators conserve energy in part through reduced liver mitochondrial oxidation in response to cold temperature (24,25), decreasing SDH activity (26,27) that in turn impairs cellular respiration as they lower body temperature and metabolic rate. In contrast, winter juvenile shrews show upregulated oxidative phosphorylation pathways compared to summer juveniles. While Dehnel’s phenomenon reduces resting metabolic rate in winter (12), shrews remain active and do not reduce their body temperature as hibernators do. Mitochondrial regulation in shrews is opposite to that of hibernators and may use known temperature-sensitive oxidative phosphorylation pathways to improve mitochondrial efficiency, enhance metabolic plasticity, or maintain internal temperatures despite the reduction in metabolic rate and body size. This hypothesis can be further tested by examining mitochondrial oxygen consumption in the liver throughout Dehnel’s phenomenon.

During autumn and winter, shrews appeared to meet their energetic needs by activating responses similar to those triggered by starvation. When blood sugar is low, mammals promote gluconeogenesis, a process in which the liver produces free glucose from non-carbohydrate substrates to avoid cellular starvation (19). This process has previously been linked to seasonal size change in shrews (10). Glucose-6-phosphatase (G6P) enzyme levels peak during autumn and decline through spring, suggesting gluconeogenesis is highest in autumn and early winter when snow is too scarce to offer insulation, increasing the need for glucose to prevent starvation. We discovered an upregulation of gluconeogenesis pathways in winter juveniles, including the overexpression of aldehyde dehydrogenases (*ALDH1B1* and *ALDH3A2*) and *G6PC1*. *G6PC1* encodes one of three subunits of G6PC, a key gluconeogenesis enzyme in the final step of gluconeogenesis, which catalyzes the formation of D-glucose that can then be released into the blood. We propose the upregulation of this gene, along with others in the pathway, regulates an increase in gluconeogenesis (10). Alongside elevated lipid metabolism and oxidative phosphorylation, these findings indicate shrews maximize all available energy resources during winter (Fig 4).

**Fig 4.**
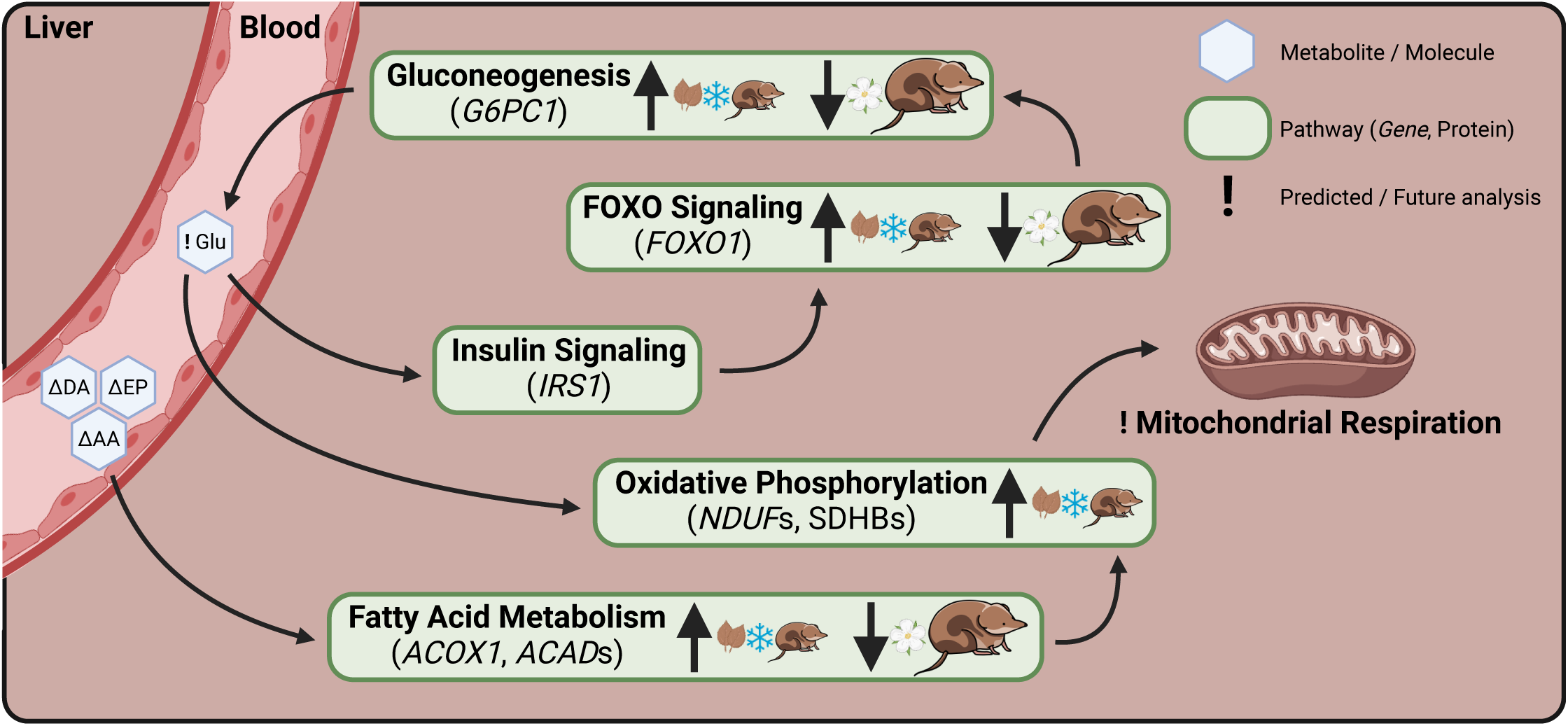
Summary of dynamic changes in shrew metabolism associated with seasonal size change, with proposed regulation of Dehnel’s cycle. Depleted lipid metabolites in the blood and increased RNA expression of fatty acid metabolism genes highlights the importance of rapid fat turnover in winter juveniles. FOXO signaling regulates gluconeogenesis, which is also upregulated in winter juveniles, providing another energy source. Both FOXO signaling and fatty acid metabolism promote changes in oxidative phosphorylation, found in both transcriptomic and proteomic analyses, and have large effects on mitochondrial respiration in the liver.

Large changes in mRNA and proteins only weakly supported one another when comparing winter juveniles to spring adults, but changes in gene networks across all seasons parallel regrowth. We identified a large network negatively correlated with liver size and enriched with genes involved in FOXO and insulin signaling. Central to this network is the forkhead box transcription factor 1 (*FOXO1*), upregulated during shrinkage and downregulated during growth. In mice, FOXO1 stimulates gluconeogenesis that promotes hepatic glucose production and release (28–30), which can subsequently impede processes like lipid metabolism in response to fasting glucose levels (29–32). In shrinking and regrowing shrews, FOXO signaling may thus regulate organismal energy homeostasis and size by simultaneously regulating both glucose and lipid use to meet shifting energy demands.

Increased FOXO signaling may, however, incur costs to shrew longevity. In model organisms including *C. elegans*, *Drosophila*, mice and humans, FOXO1 overexpression can reduce size and extend lifespan, while inhibition leads to aging and senescence (33–36). Compared to other mammals, *S. araneus* evolved Dehnel’s phenomenon, a rare size plasticity (1–3), as well as extreme and disproportionately high metabolic rate (37) and short lifespan (38). FOXO1 cycling could explain this unique combination of traits. Should findings from model organisms apply in the shrew, FOXO1-dependent glucose utilization regulates shrinking reducing mortality during harsh winters but undermines shrew longevity when reversed in spring as a terminal investment for growth and reproduction. Genes within this network are therefore key candidates for exploring both the regulatory mechanisms of size change in Dehnel’s phenomenon and metabolic change correlated with lifespan in mammals.

Despite valuable insights, there are limitations and questions outside the scope of this study. Metabolic and regulatory change was extensively characterized throughout seasonal size changes, but functional validation of current hypotheses is needed. For example, future studies could analyze mitochondrial respiration to detect changes in seasonal efficiency or probe blood glucose levels. Seasonal size change occurs in other shrew organs, and several mechanisms proposed here involve interorgan communication, including communication between the hypothalamus and liver through the endocrine system. Thus, it is important to investigate other tissues, such as adipocytes in response to changes in lipid turnover, or the brain, which is rich in fats and experiences extraordinary size plasticity, particularly in the hippocampus and cortex. Additionally, we could investigate proposed regulatory mechanisms of longevity by examining if shrews that do not undergo Dehnel’s phenomenon, and if, lacking FOXO signaling, they exhibit extended lifespans. Such future tests could reveal insights into natural neurodegenerative processes, their links to metabolic dysfunction and longevity, and provide targets for potential treatments for aging and neurodegeneration.

### Conclusion

Dehnel’s phenomenon was known to decrease energy spent on large and energy expensive tissue (8,13,37,39), but associated metabolic and regulatory shifts were unknown. Analyzing changes to the shrew liver and blood plasma, we discovered a decrease in polyunsaturated fatty acid metabolites and altered oxidative phosphorylation pathways, potentially related to mitochondrial respiration, alongside upregulated lipid and glucose metabolism during winter shrinkage. These changes distinguish Dehnel’s phenomenon from other wintering strategies. As in hibernation, Dehnel’s phenomenon alters mitochondrial function and relies upon increased lipid metabolism to meet winter energetic demands, but rapidly depletes fats, supplementing energetic demands with gluconeogenesis. These metabolic fluctuations and subsequent size change seem to be regulated by FOXO and insulin signaling pathways that regulate both size and lifespan in several model organisms (33–36). Along with key genes (*ACOX1*, *SDHB*, NDUFs), these pathways are strong candidate regulators for Dehnel’s phenomenon and link metabolism, size, and lifespan in mammals.

## Material and Methods

### Sample Collection and RNA Sequencing

Eurasian common shrews were trapped using insulated wooden traps in Radolfzell, Germany, at five different stages of development: large-brained summer juveniles (n=5), shrinking autumn juveniles (n=4), small winter juveniles (n=5), regrowing spring adults (n=5), and regrown summer adults (n=5). Shrews were aged and sexed based on tooth wear, fur appearance, and gonad development prior to euthanasia. Fewer females than males were sampled in juvenile stages (n=6 females). At that age, shrews are prepubescent, not sexually dimorphic, and only minor sex dimorphism was detected in the size change of representative brain regions in a previous study (Lázaro et al. 2018). Female under sampling is unlikely to influence gene expression, but sex was used as a covariate in subsequent models, accounting for this skew.

Blood was collected prior to perfusion through the heart, let stand for 15 minutes, spun down at 1200 rpm, the serum pipetted off and then stored at −80°C. Shrews were perfused under anesthesia via the vascular system with PAXgene Tissue Fixative. Livers were removed and weighed to confirm the occurrence of DP and then placed immediately into PAXgene Tissue Stabilizer. All samples were placed in stabilizer (2-24 hours after extraction) and then stored in liquid nitrogen. RNA was extracted with a modified Qiagen Micro RNAeasy protocol used for small amounts of mammalian tissue (40). RNA was sent to Genewiz for quality control, library preparation, and sequencing. Quantity of RNA was measured with a Nanodrop spectrophotometer and quality of RNA assessed with RNA ScreenTape. Libraries were prepared with standard polyA selection and sequenced as 150bp pair-end reads for a target of 15-25 million reads per sample. Body and liver weight were tested for significant change in mass using a t-test between seasons.

### Metabolomics

Metabolite analysis was carried out by MS-Omics using a Thermo Scientific Vanquish LC (UPLC) coupled to Thermo Q Exactive HF MS. Identification of compounds were performed at four levels; Level 1: identification by retention times (compared against in-house authentic standards), accurate mass (with an accepted deviation of 3ppm), and MS/MS spectra, Level 2a: identification by retention times (compared against in-house authentic standards), accurate mass (with an accepted deviation of 3 ppm). Level 2b: identification by accurate mass (with an accepted deviation of 3ppm), and MS/MS spectra. MetaboAnalyst was used to test for changes in concentration between seasons (41). Concentrations for each sample were normalized by the average concentration of summer adult samples, log transformed and auto scaled. A one-way ANOVA was used to test for any significant seasonal difference in metabolite concentration, with p-values corrected with a false discovery rate of 0.05. Within each significant metabolite, t-tests were used to identify pairwise differences in mean between each season. Samples were hierarchically clustered using significant genes and visualized using a heatmap.

### Differential Gene Expression Analysis

Tests for differential gene expression were conducted to identify genes whose expression changes during Dehnel’s phenomenon. We trimmed adapters and filtered reads using fastp (42) and removed samples with extreme read counts (∼10-fold). Gene counts were quantified by pseudo aligning to the 19,296 protein coding genes of the *S. araneus* genome (sorAra2; GCF_000181275.1) using Kallisto (43). Counts were normalized for each tissue using DESeq2 median of ratios (44). In the liver, summer juveniles and winter juveniles were compared to infer regulatory processes underpinning shrinkage, while winter juveniles were tested against spring adults to explore processes underlying regrowth. Normalized counts for each gene were fitted with a negative binomial generalized linear model and then we used a Wald test to examine the statistical association between the log-fold change and the season parameter, with p-values corrected using the Benjamini and Hochberg procedure (45). We then identified significantly enriched pathways with differentially expressed genes using a ranked gene set enrichment of KEGG pathways using fgsea (46).

### Weighted Gene Co-expression Network Analysis

We used the WGCNA package (47) package to create co-expression networks for the liver and hippocampus and determined if these correlations between genes are significantly correlated with our shrinkage and regrowth phenotypes. We calculated the Pearson correlation for expression between each gene pair, which was then transformed to an adjacency matrix using a soft-thresholded power adjacency function described in Langfelder and Horvath (47). The power function, which reduced the spurious connectivity between genes from noise by increasing sensitivity, was determined by maximizing the scale-free topology criterion (maximizing model fit while saturating mean connectivity). The resulting adjacency matrix was then used to calculate a topological overlap matrix (dissimilarity matrix), in which dissimilarity distances grouped genes into modules using fuzzy cmeans hierarchical clustering. We removed any module with fewer than 25 genes, and merged gene modules that were found to have similar eigengene values. Then, eigengene significance between the correlation of liver size to each module’s eigengene value was calculated. Cytoscape (48) was used to visualize significant networks and calculate network statistics such as node (gene) connectivity. Gene set enrichment of each module was analyzed using DAVID Gene Functional Classification Tool to test for GO enrichment (49).

### Proteomics

Liver tissue from *Sorex araneus* was isolated and protein homogenates were prepared by transferring the liver tissue to a dissection buffer containing 10% sucrose (VWR), imidazole (Merck Millipore), EDTA (Sigma-Aldrich), Pefabloc (Sigma-Aldrich) and leupeptin (VWR). The mixture was homogenized (T10 basic ULTRA-TURAX homogenizer IKA®) for 20 sec; then, samples were centrifuged at 1000 g for 15 min. Supernatants were stored at −20°C. Proteomic sample preparation was performed using the PreOmics, Cat. No.: 00027 iST 96x kit. The samples were vacuum centrifuged overnight, and the dry peptide product was stored at −80°C until analysis.

Peptides were resuspended in a solution containing 2% acetonitrile (ACN), 0.1% formic acid (FA) and 0.1% TFA, and then, peptides were briefly sonicated. Five micrograms of total peptide material were analyzed per liquid chromatography–mass spectrometry analysis. Samples were analyzed using a UPLC-nanoESI MS/MS setup with a NanoRSLC system (Dionex, Sunnyvale, CA, USA). The system was coupled online with an emitter for nanospray ionization (New Objective PicoTip 360-20-10) to a Q Exactive HF mass spectrometer (Thermo Scientific, Waltham, USA). The peptide material was loaded onto a 2-cm trapping reversed-phase Acclaim PepMap RSLC C18 column (Dionex) and separated using an analytical 75-cm reversed-phase Acclaim PepMap RSLC C18 column (Dionex). Both columns were kept at 60°C. The sample was eluted with a gradient of 90% solvent A (0.1% FA, 0.1% TFA) and 10% solvent B (0.1% FA, 0.1% TFA in ACN), which was increased to 7% solvent B on a 1-min ramp gradient at a constant flow rate of 300 nL/min. Subsequently, the gradient was raised to 30% solvent B on a 45-min ramp gradient. The mass spectrometer was operated in positive mode, selecting up to 20 precursor ions with a mass window of m/z 1.6 based on highest intensity for higher-energy collisional dissociation (HCD) fragmenting at a normalized collision energy of 27. Selected precursors were dynamically excluded for fragmentation for 30 s.

A label-free relative quantitation analysis was performed using MaxQuant 1.5.7.4 software (50). Raw files were searched against the *S. araneus* genome (GCF_027595985.1_mSorAra2.pri_genomic.gtf). All standard settings were employed with carbamidomethylation (C) as a static peptide modification and deamidation (NQ), oxidation (M), formylation (N-terminal and K), and protein acetylation (N-terminal) as variable modifications. The output contained a list of proteins identified <1% false discovery rate, and their abundances were further filtered and processed using the Perseus v1.5.6.0 platform (51). All reverse hits that identified proteins were removed from further analysis, and the data were log2-transformed to approximate a normal distribution. Two or more unique peptides were required for protein quantitation. Additionally, a non-zero quantitation value in at least 70% of the samples in one of the groups was required for the quantifiable proteins.

We inferred potential functional changes associated with differences in protein concentrations and analyzed correlations with transcriptomic data. We identified significantly enriched pathways of protein using a ranked gene set enrichment of KEGG pathways using fgsea (46). Additionally, to investigate the relationship between seasonal changes in protein concentration and RNA abundance, we performed a linear regression analysis. Differences in protein concentration were modeled as a function of the corresponding gene expression change using the lm function in R.

## Acknowledgments

We thank Joshua Rest, Krishna Veeramah, and Tanya Lama who have provided helpful feedback on initial results and previous versions of the manuscript. We thank Michal Oklinski for help with initial dissections.

## Supporting Information

### Data Availability

Further details of experimental protocols are described in the Expanded View. Data openly available on Dryad https://doi.org/10.5061/dryad.pc866t1w3 with reproducible code found on GitHub: https://github.com/wrthomas315/Dehnels_Seasonal_RNAseq2/tree/main/data. Primary sequencing was deposited on the NCBI SRA, BioprojectPRJNA941271.

### Author Contributions and Funding Disclosure

LD, DD, AC, and JN conceived and funded project. CB, MM, and DD collected, measured, and sampled shrews. WRT designed and conducted genomic analyses. WRT, JHJ, JN, and AC analyzed metabolite concentrations. WRT visualized data and wrote initial draft. LD, DD, JN, DC, and AC contributed to review and editing of draft. All authors contributed to data interpretation. WRT and research were supported by the Human Frontiers in Science Program Award (RGP0013/2019) to DD, LMD, and JN, and by the Stony Brook University Presidential Innovation and Excellence Fund to LMD.

## References

1. Dehnel A. Studies of the genus Sorex L. Ann Univ M Curie-Sklod. 1949;4(2):17–104.

2. Pucek Z. Seasonal and age changes in the weight of internal organs of shrews. Acta Theriol (Warsz). 1965;10:369–438.

3. Lázaro J, Hertel M, Sherwood CC, Muturi M, Dechmann DKN. Profound seasonal changes in brain size and architecture in the common shrew. Brain Struct Funct [Internet]. 2018;223(6):2823–40. Available from: 10.1007/s00429-018-1666-5

4. Lázaro J, Dechmann DKN. Dehnel’s phenomenon. Curr Biol. 2021;31(10):R463–5.

5. Pucek M. Water contents and seasonal changes of the brain-weight in shrews. Acta Theriol (Warsz). 1965;10(1955):353–67.

6. Ray S, Li M, Koch SP, Mueller S, Boehm-Sturm P, Wang H, et al. Seasonal plasticity in the adult somatosensory cortex. Proc Natl Acad Sci U S A. 2020;117(50):32136–44.

7. Genoud M, Isler K, Martin RD. Comparative analyses of basal rate of metabolism in mammalslJ: data selection does matter. Biol Rev. 2018;93:404–38.

8. Keicher L, O’Mara MT, Voigt CC, Dechmann DKN. Stable carbon isotopes in breath reveal fast metabolic incorporation rates and seasonally variable but rapid fat turnover in the common shrew (Sorex araneus). J Exp Biol. 2017;220(15):2834–41.

9. Churchfield S, Rychlik L, Taylor JRE. Food resources and foraging habits of the common shrew, Sorex araneus: Does winter food shortage explain Dehnel’s phenomenon? Oikos. 2012;121(10):1593–602.

10. Hyvärinen H. Wintering strategy of voles and shrews in Findland. Winter Ecol Small Mamm. 1984;10:139–48.

11. Lázaro J, Hertel M, Muturi M, Dechmann DKN. Seasonal reversible size changes in the braincase and mass of common shrews are flexibly modified by environmental conditions. Sci Rep. 2019;9(1):1–10.

12. Taylor JRE, Rychlik L, Churchfield S. Winter reduction in body mass in a very small, nonhibernating mammal: Consequences for heat loss and metabolic rates. Physiol Biochem Zool. 2013;86(1):9–18.

13. Schaeffer PJ, O’Mara MT, Breiholz J, Keicher L, Lázaro J, Muturi M, et al. Metabolic rate in common shrews is unaffected by temperature, leading to lower energetic costs through seasonal size reduction. R Soc Open Sci. 2020;7(4).

14. Auteri GG. A conceptual framework to integrate cold-survival strategies: Torpor, resistance and seasonal migration. Biol Lett. 2022;18(5):1–6.

15. Boyer BB, Barnes BM. Molecular and metabolic aspects of mammalian hibernation. Bioscience. 1999;49(9):713–24.

16. Chazarin B, Storey KB, Ziemianin A, Chanon S, Plumel M, Chery I, et al. Metabolic reprogramming involving glycolysis in the hibernating brown bear skeletal muscle. Front Zool. 2019;16(1):1–21.

17. Faherty SL, Luis Villanueva-Cañas J, Klopfer PH, Albà MM, Yoder AD. Gene expression profiling in the hibernating primate, Cheirogaleus medius. Genome Biol Evol. 2016;8(8):2413–26.

18. Villanueva-Cañas JL, Faherty SL, Yoder AD, Albà MM. Comparative genomics of mammalian hibernators using gene networks. Integr Comp Biol. 2014;54(3):452–62.

19. Rui L. Energy Metabolism in the Liver Liangyou Rui. Physiol Behav. 2014;176(5):139– 48.

20. Lu D, He A, Tan M, Mrad M, El Daibani A, Hu D, et al. Liver ACOX1 regulates levels of circulating lipids that promote metabolic health through adipose remodeling. Nat Commun. 2024;15(1).

21. Elaine Epperson L, Karimpour-Fard A, Hunter LE, Martin SL. Metabolic cycles in a circannual hibernator. Physiol Genomics. 2011;43(13):799–807.

22. Drew KL, Rice ME, Kuhn TB, Smith MA. Neuroprotective adaptations in hibernation: Therapeutic implications for ischemia-reperfusion, traumatic brain injury and neurodegenerative diseases. Free Radic Biol Med. 2001;31(5):563–73.

23. Rice SA, Mikes M, Bibus D, Berdyshev E, Reisz JA, Gehrke S, et al. Omega 3 fatty acids stimulate thermogenesis during torpor in the Arctic Ground Squirrel. Sci Rep [Internet]. 2021;11(1):1–14. Available from: 10.1038/s41598-020-78763-8

24. Chaffee RR, Hoch FL, Lyman CP. Mitochondrial oxidative enzymes and phosphorylations in cold exposure and hibernation. Am J Physiol. 1961;201(1):29–32.

25. Staples JF, Brown JCL. Mitochondrial metabolism in hibernation and daily torpor: A review. J Comp Physiol B Biochem Syst Environ Physiol. 2008;178(7):811–27.

26. Armstrong C, Staples JF. The role of succinate dehydrogenase and oxaloacetate in metabolic suppression during hibernation and arousal. J Comp Physiol B Biochem Syst Environ Physiol. 2010;180(5):775–83.

27. Gehnrich SC, Aprille JR. Hepatic gluconeogenesis and mitochondrial function during hibernation. Comp Biochem Physiol -- Part B Biochem. 1988;91(1):11–6.

28. Matsumoto M, Pocai A, Rossetti L, DePinho RA, Accili D. Impaired Regulation of Hepatic Glucose Production in Mice Lacking the Forkhead Transcription Factor Foxo1 in Liver. Cell Metab. 2007;6(3):208–16.

29. Xiong X, Tao R, DePinho RA, Dong XC. Deletion of Hepatic FoxO1/3/4 Genes in Mice Significantly Impacts on Glucose Metabolism through Downregulation of Gluconeogenesis and Upregulation of Glycolysis. PLoS One. 2013;8(8):1–11.

30. Zhang W, Patil S, Chauhan B, Guo S, Powell DR, Le J, et al. FoxO1 regulates multiple metabolic pathways in the liver effects on gluconeogenic, glycolytic, and lipogenic gene expression. J Biol Chem [Internet]. 2006;281(15):10105–17. Available from: 10.1074/jbc.M600272200

31. Tikhanovich I, Cox J, Weinman SA. Forkhead box class O transcription factors in liver function and disease. J Gastroenterol Hepatol. 2013;28(1):125–31.

32. Yang Z, Roth K, Agarwal M, Liu W, Petriello MC. The transcription factors CREBH, PPARa, and FOXO1 as critical hepatic mediators of diet-induced metabolic dysregulation. J Nutr Biochem [Internet]. 2021;95:108633. Available from: 10.1016/j.jnutbio.2021.108633

33. Gami MS, Wolkow CA. Studies of Caenorhabditis elegans DAF-2/insulin signaling reveal targets for pharmacological manipulation of lifespan. Aging Cell. 2006;5(1):31–7.

34. Hwangbo DS, Garsham B, Tu MP, Palmer M, Tatar M. Drosophila dFOXO controls lifespan and regulates insulin signalling in brain and fat body. Nature. 2004;429(6991):562–6.

35. Katic M, Kahn CR. The role of insulin and IGF-1 signaling in longevity. Cell Mol Life Sci. 2005;62(3):320–43.

36. Satoh A, Brace CS, Rensing N, Cliften P, Wozniak DF, Herzog ED, et al. Sirt1 extends life span and delays aging in mice through the regulation of Nk2 Homeobox 1 in the DMH and LH. Cell Metab [Internet]. 2013;18(3):416–30. Available from: 10.1016/j.cmet.2013.07.013

37. Taylor JRE. Evolution of Energetic Strategies in Shrews. In: Evolution of Shrews. 1998. p. 309–46.

38. Healy K, Guillerme T, Finlay S, Kane A, Kelly SBA, McClean D, et al. Ecology and mode-of-life explain lifespan variation in birds and mammals. Proc R Soc B. 2014;281.

39. Pucek Z. Seasonal and age change in shrews as an adaptive process. Symp zool Soc Lond. 1970;26:189–207.

40. Yohe LR, Davies KTJ, Simmons NB, Sears KE, Dumont ER, Rossiter SJ, et al. Evaluating the performance of targeted sequence capture, RNA-Seq, and degenerate-primer PCR cloning for sequencing the largest mammalian multigene family. Mol Ecol Resour. 2020;20(1):140–53.

41. Pang Z, Chong J, Zhou G, De Lima Morais DA, Chang L, Barrette M, et al. MetaboAnalyst 5.0: Narrowing the gap between raw spectra and functional insights. Nucleic Acids Res. 2021;49:388–96.

42. Chen S, Zhou Y, Chen Y, Gu J. Fastp: An ultra-fast all-in-one FASTQ preprocessor. Bioinformatics. 2018;34(17):i884–90.

43. Bray NL, Pimentel H, Melsted P, Pachter L. Near-optimal probabilistic RNA-seq quantification. Nat Biotechnol. 2016;34(5):525–7.

44. Love MI, Huber W, Anders S. Moderated estimation of fold change and dispersion for RNA-seq data with DESeq2. Genome Biol. 2014;15(550):1–21.

45. Benjamini Y, Hochberg Y. Controlling the False Discovery RatelJ: A Practical and Powerful Approach to Multiple Testing. J R Stat Soc. 1995;57(1):289–300.

46. Korotkevich G, Sukhov V, Budin N, Atryomov MN, Sergushichev A. Fast gene set enrichment analysis. bioRxiv. bioRxiv. 2021;

47. Langfelder P, Horvath S. WGCNA: An R package for weighted correlation network analysis. BMC Bioinformatics. 2008;9.

48. Shannon P, Markiel A, Ozier O, Nitin SB, Wang JT, Ramage D, et al. Cytoscape: A Software Environment for Integrated Models. Genome Res [Internet]. 2003;13(22):2498–504. Available from: http://ci.nii.ac.jp/naid/110001910481/

49. Huang DW, Sherman BT, Lempicki RA. Systematic and integrative analysis of large gene lists using DAVID bioinformatics resources. Nat Protoc. 2009;4(1):44–57.

50. Tyanova S, Temu T, Cox J. The MaxQuant computational platform for mass spectrometry-based shotgun proteomics. Nat Protoc [Internet]. 2016;11(12):2301–19. Available from: 10.1038/nprot.2016.136

51. Tyanova S, Temu T, Sinitcyn P, Carlson A, Hein MY, Geiger T, et al. The Perseus computational platform for comprehensive analysis of (prote)omics data. Nat Methods. 2016;13(9):731–40.

